# Indirect pathway of caudate tail for choosing good objects in periphery

**DOI:** 10.1101/547232

**Authors:** Hidetoshi Amita, Okihide Hikosaka

## Abstract

Choosing good objects is essential for real life, which is controlled mainly by the basal ganglia. For that, a subject need to not only find good objects, but ‘reject’ bad objects. To reveal this ‘rejection’ mechanism, we created a sequential saccade choice task for monkeys and studied the indirect pathway of caudate tail mediated by cvGPe (caudal-ventral globus pallidus externus). The inhibitory responses of cvGPe neurons to bad objects were smaller when the monkey made saccades to them by mistake. Moreover, experimental reduction of the inhibitory response by local injection of bicuculline (GABA_A_ antagonist) disabled the monkey to reject bad objects. In conclusion, rejecting bad objects is crucial for goal-directed behavior, which is controlled by the indirect pathway in the basal ganglia.

## Introduction

Experts as chess masters can rapidly and automatically perceive visual information and make decisions based on skilled memory(*1, 2*). Previous studies suggest that the basal ganglia are involved in these skills in humans and monkeys(*3–6*), but their neuronal mechanisms are unclear. The basal ganglia dysfunctions cause various cognitive and movement disorders(*7, 8*), and variable models had been proposed for explaining their mechanisms(*9–17*). Notably, it has been still discussed on how two main circuits (i.e., direct pathway and indirect pathway) cooperate or compete in the basal ganglia. While the direct pathway facilitates action (Go) and the indirect pathway suppresses action (No-go)(*18, 19*), both the direct and indirect pathways are active during action initiation(*20–22*). To investigate the neuronal mechanisms in the basal ganglia for skills, we focused on the posterior basal ganglia circuits.

The posterior basal ganglia circuits originating from the caudate tail (CDt) send signals to the superior colliculus (SC) through the caudal-dorsal-lateral substantia nigra pars reticulata (cdlSNr) and/or the caudal-ventral globus pallidus externus (cvGPe)(*23–25*). These areas have two key features: high-capacity/long-term value memory and peripheral vision. Many neurons along the CDt-circuits (CDt, cvGPe, cdlSNr and SC) sustain their value preference for many objects over a period of months, and discriminate between good and bad objects in peripheral receptive fields (mostly in the contralateral hemisphere) regardless of eccentricity(*23, 24, 26, 27*). However, their roles in active choice is still unclear, because these neuronal data were based on passive conditions. Here, we investigated whether the CDt-circuits are involved in value-based action/object choice, and how the indirect pathway works in cooperation with the direct pathway.

## Results

Previous studies revealed that the neurons encoding value of objects are localized in cvGPe and cdlSNr(*23, 24*). We first examined these neurons using the object-reward association learning task (Fig. 1A) and the passive viewing task (Fig. 1B). Most neurons in cvGPe and cdlSNr responded differently to good and bad objects (examples shown in Fig. 1C and D), consistent with previous studies. Bad objects inhibited cvGPe neurons (Fig. 1C) but excited cdlSNr neurons (Fig. 1D); good objects caused opposite responses. These reversed response patterns between cvGPe and cdlSNr, especially to bad objects, can be explained by sequential inhibitory connections (Fig. 1E): 1) from CDt to cvGPe, 2) from cvGPe to cdlSNr, 3) from cdlSNr to SC (i.e., indirect pathway). These results suggest that saccades are suppressed by the indirect pathway (Fig. 1E).

**Fig. 1.**
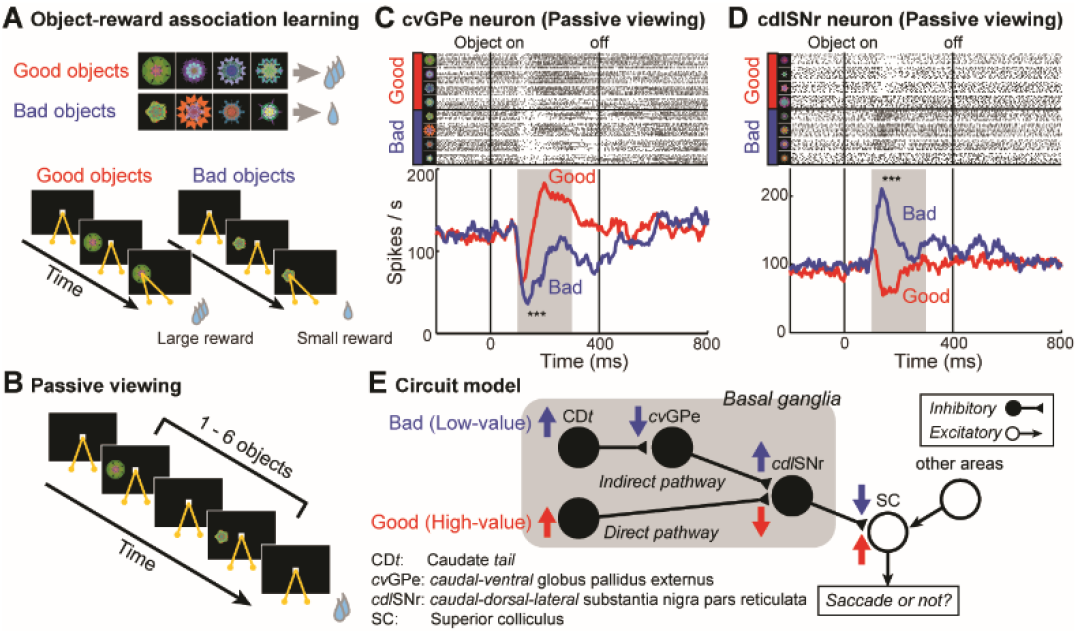
Value-coding circuits in the basal ganglia. (A) Object-reward association learning task. (B) Passive viewing task. (C) Representative cvGPe neuron inhibited by bad objects during passive viewing task (*n* = 33 trials, t (32) = 5.17, \*\*\**P* < 0.001, Paired *t*-test). Its activity, which is shown by raster plots (above) and spike density functions (below), is aligned on object onset and offset. Responses are shown separately for all eight objects (above) and as the averaged responses to good and bad objects (below). (D) Representative cdlSNr neuron excited by bad objects (*n* = 34 trials, t (33) = −8.38, \*\*\**P* < 0.001, Paired *t*-test). (E) Neuronal circuit model for value-based saccades. Value signals for bad objects (blue) are sent through the indirect pathway (CDt-cvGPe-cdlSNr), while value signals for good objects (red) are sent through the direct pathway (CDt-cdlSNr), both within the basal ganglia (gray region). Upward and downward arrows indicate excitation and inhibition, respectively.

We then tested the above hypothesis by using a new procedure called ‘sequential saccade choice task’ (Fig. 2A, Movie S1 and S2). In each block of trials, 8 fractal objects (4 good, 4 bad) appeared sequentially and randomly at peripheral positions, until the subject chose one object. In this case (Fig. 2A and Movie S1), the subject first rejected two bad objects (1st object: no saccade, 2nd object: saccade to object, immediately followed by another saccade back to center) and then accepted a good object (saccade to object, followed by maintained gaze) which provided a reward. If the subject accepted a bad object, no reward was given. The subject experienced multiple sets of objects (4 sets for ZB, 4 for SP) using this task. In the first learning session (when all objects were new), the subject made a saccade to most good as well as bad objects (Fig. 2B, left). Across the subsequent sessions, saccade rate to bad objects gradually decreased (i.e., rejection of bad objects increased) (Fig. 2B). Bad objects were also rejected sometimes by a returning saccade (e.g., response to the 2nd object in Fig. 2A and Movie S1). Acceptance of good objects remained nearly 100 % (red curves in Fig. 2B). Importantly, the good-bad discrimination was completely maintained more than 1 month after the final learning session (Fig. 2B, right). After long-term learning (10 sessions) we recorded the activity of cvGPe and cdlSNr neurons. In response to bad objects, cvGPe neurons were significantly inhibited (Fig. 2C), while cdlSNr neurons were significantly excited (Fig. 2D). The results were similar to those during passive viewing task (Fig. 1C and D). Notably, the inhibition of cvGPe neurons by bad object was stronger when no saccade was made to the bad object (blue) than when a saccade was made (cyan) (Fig. 2C). Correspondingly, cdlSNr neurons were more strongly excited when no saccade was made (blue) than when a saccade was made (cyan) (Fig. 2D). The analysis window was set at 100 - 200 ms after object onset, because more than 80 percent of all trials showed saccade latency within 100 – 200 ms after object onset in both subjects (Fig. S1A, ZB; 83.1%, Fig. S1B, SP; 88.5%). Equivalent data are shown in more detailed for example eye trajectory (Fig. S2A), example neuronal data in cvGPe (Fig. S2B) and cdlSNr (Fig. S2C). These results suggest that the inhibition of cvGPe neurons by a bad object leads to the excitation of cdlSNr neurons, which can suppress the saccade to the bad object if the activity of cdlSNr neurons reaches a threshold (blue in Fig. 2E).

**Fig. 2.**
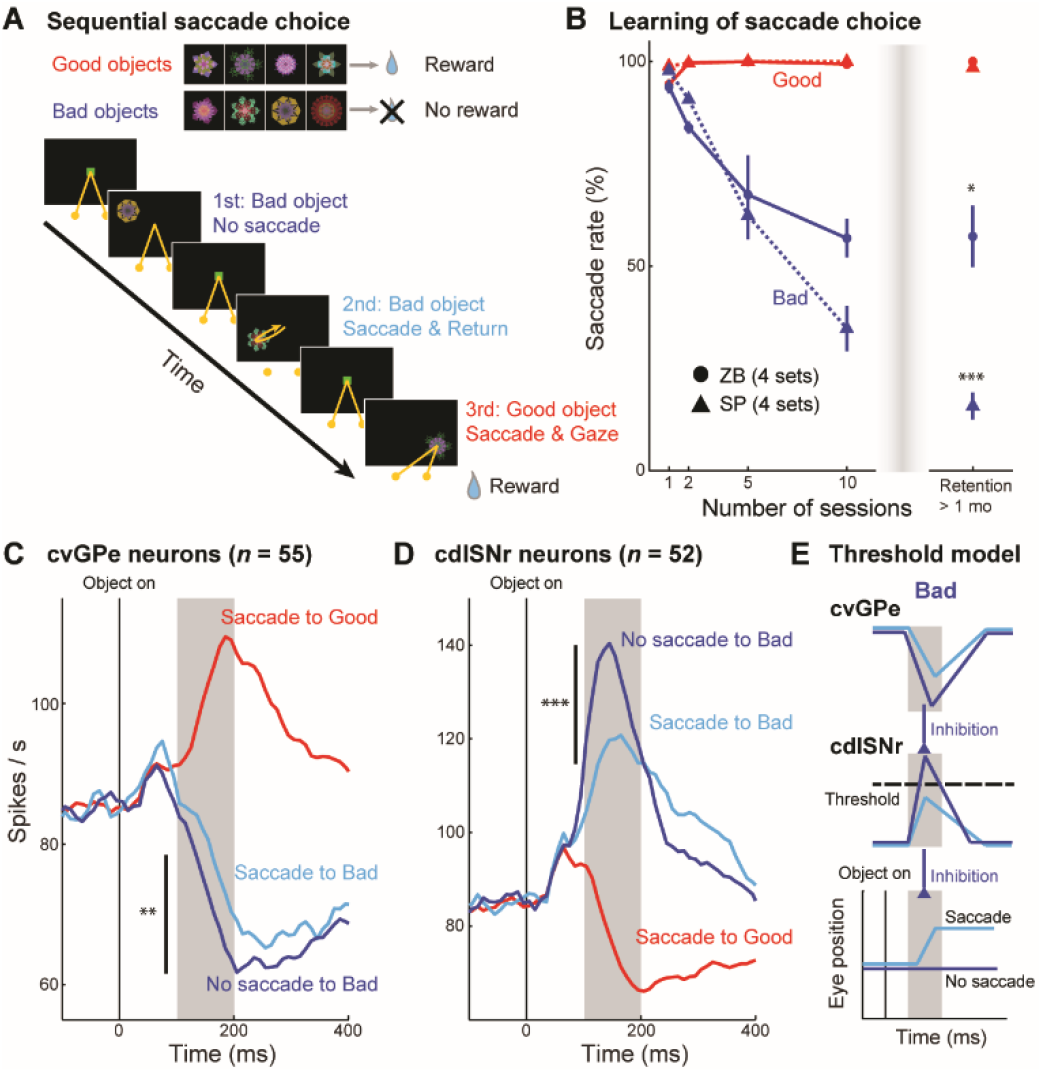
Neuronal activity for rejecting bad object. (A) Sequential saccade choice task. (B) Learning of saccade choice for each object set, averaged across 4 sets (32 objects) in each monkey (ZB and SP). Red and blue symbols indicate averaged saccade rate (%) to good and bad objects, respectively, across learning sessions. Error bars indicate S.E.M. The same task was also tested long time after the learning (Retention, > 1 month) (ZB; t (3) = - 3.87, \**P* = 0.030, Paired *t*-test, SP; t (3) = −19.4, \*\*\**P* < 0.001, Paired *t*-test). (C) Averaged activity of cvGPe neurons (*n* = 55 neurons) in response to good object (red) and bad objects (cyan and blue). The inhibition by bad objects (t (54) = 3.77, \*\*\**P* < 0.001, Paired *t*-test) was stronger when no saccade occurred (blue) than when saccade occurred (cyan) (gray period, t (54) = −2.94, \*\**P* = 0.0047, Paired *i*-test). (D) Averaged activity of cdlSNr neurons (*n* = 52 neurons), shown in the same format. The excitation by bad objects (t (51) = −7.12, \*\*\**P* < 0.001, Paired *t*-test) was stronger when no saccade occurred (blue) than when saccade occurred (cyan) (gray period, t (51) = 4.65, \*\*\**P* < 0.001, Paired *t*-test). (E) Hypothetical role of cvGPe-cdlSNr pathway to control saccade to bad objects. Blue and cyan indicate no saccade and saccade conditions, respectively. Black dashed line indicates a threshold level for suppressing a saccade.

The results so far suggest that the cvGPe-cdlSNr pathway can suppress saccades based on object values. To test this hypothesis, we experimentally manipulated the cvGPe-cdlSNr pathway by injecting bicuculline (GABA_A_ antagonist) locally into cvGPe (Fig. 3A and B). We chose bicuculline because cvGPe neurons receive massive GABAergic inputs, mainly from CDt(25) (Fig. 1E). We also recorded the activity of single cdlSNr neuron continuously before and after the injection (Fig. 3A), while the subject continued to perform the sequential saccade choice task (Fig. 2A). We found that bicuculline injection changed both saccade behavior (Fig. 3C and Movie S3) and cdlSNr neuronal activity (Fig. 3D) during the sequential saccade choice task. The probability of saccades to bad objects increased significantly from about 25% to almost 95% (blue bars in Fig. 3C). In other words, the subjects were no longer able to suppress saccades to bad objects when GABAergic inputs to cvGPe neurons were blocked. This occurred for bad objects located only contralateral to the bicuculline injection side (Fig. S3A). Injection of saline in cvGPe led to no significant change (Fig. S3B and Movie S4). Moreover, the excitatory response of the cdlSNr neuron to bad objects decreased significantly (Fig. 3D, also see Fig. 2D as control data). These data support the schematic model we proposed (Fig. 3E compared with Fig. 2E). In the current experiment, we have shown that the experimental manipulation of neuronal activity (by bicuculline) changed saccade behavior, as the model predicts. In detail (as shown in Fig. 3E), bicuculline suppressed the inhibitory response of cvGPe neurons to bad objects, which led to the suppression of the excitatory response of cdlSNr neurons (blue dotted line shown in Fig. 3D). The subject was then unable to suppress saccades to bad objects because the inhibition of SC saccadic neurons by cdlSNr neurons was too weak (i.e., below threshold). Notably, the probability of saccades to good objects remained 100% (red bars in Fig. 3C) and the inhibition of cdlSNr neurons in response to good objects did not significantly change after the bicuculline injection (red dotted line in Fig. 3D).

**Fig. 3.**
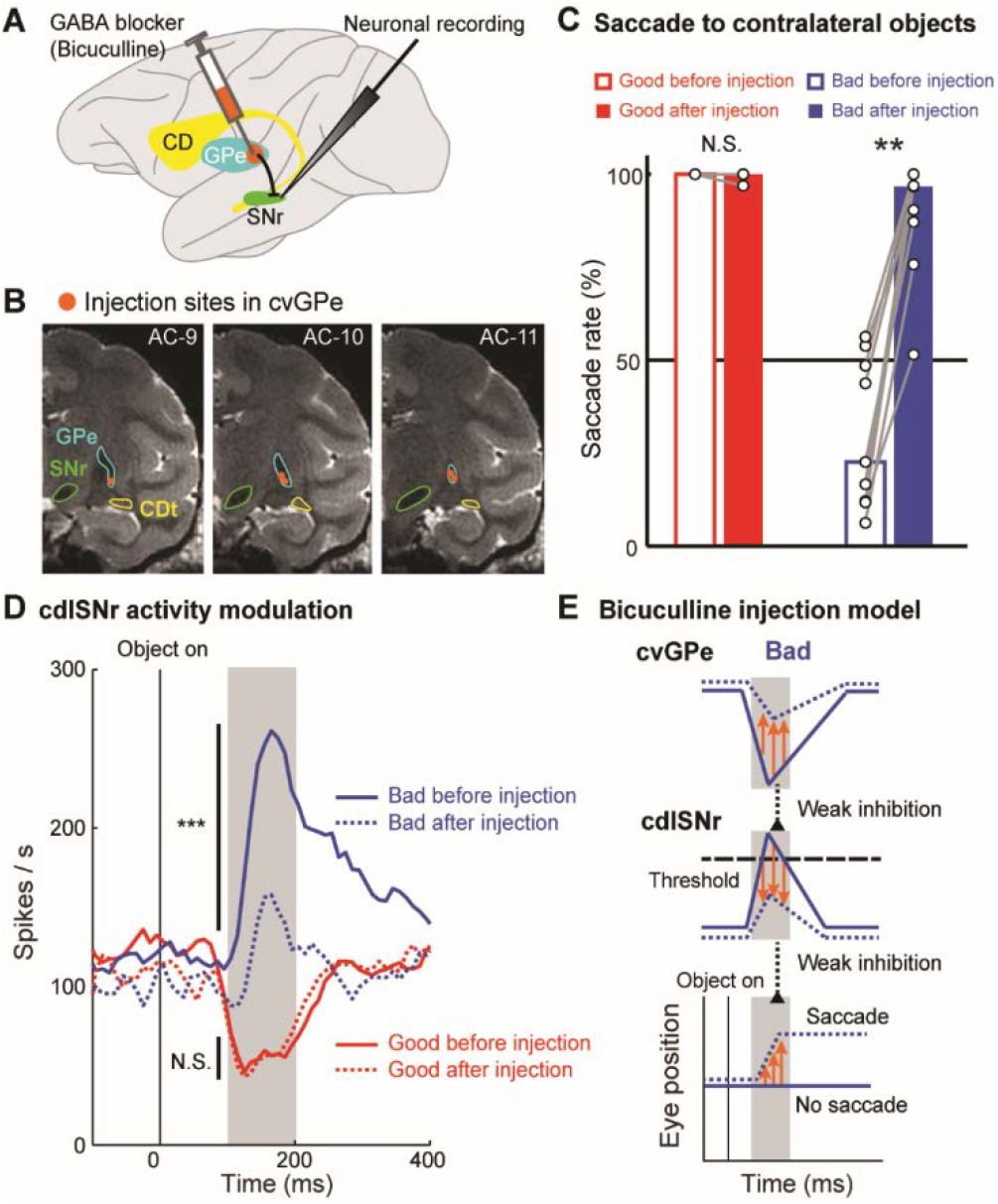
Blocking of the indirect pathway impairs rejection of bad object during sequential saccade choice task. (A) Local injection of GABA blocker (bicuculline) in cvGPe, while neuronal activity in cdlSNr was recorded. (B) MR images showing five injection sites (orange) in cvGPe of monkey ZB that were located in three coronal sections (9, 10, and 11 mm posterior to the anterior commissure). (C) Comparison of saccade rates between before and after the bicuculline injection in response to contralateral good objects (red; *n* = 9 sessions, *P* = 1.00, Wilcoxon signed-rank test) and bad objects (blue; *n* = 9 sessions, \*\**P* = 0.0039, Wilcoxon signed-rank test). Each bar indicates median of saccade rate to contralateral objects. Each pair of connected dots depicts a single injection session. (D) Comparison of one cdlSNr neuron’s activity between before and after the bicuculline injection in response to good objects (red; *n* = 64 trials, *P* = 0.99, Mann-Whitney *U* test) and bad objects (blue; *n* = 64 trials, \*\*\**P* < 0.001, Mann-Whitney *U* test). (E) Schematic model for explaining the neuronal and behavioral effects (orange arrows) of GABA blocker in cvGPe. Blue solid and dotted lines indicate responses to bad objects before and after the bicuculline injection.

We also examined the effect of bicuculline in cvGPe on the choice of objects using the simultaneous saccade choice task (Fig. 4A), instead of the sequential saccade choice task (Fig. 2A). In the simultaneous saccade choice task, a pair of good and bad objects was presented in either the contralateral or ipsilateral hemifield randomly. During each saccade, the objects were replaced by white dot. The outcome (i.e., large or small reward) was based on the choice (Fig. 4A, below). The choice rate of good objects in the contralateral hemifield significantly decreased after the bicuculline injection (Fig. 4B). In other words, the subject’s choice of bad objects increased to the chance level (50%). This did not occur for objects located ipsilateral to the bicuculline injection side (Fig. S4A, right). Injection of saline in cvGPe led to no significant change (Fig. S4B). These data together demonstrated that the cvGPe-cdlSNr pathway suppresses saccades to bad objects, which enables the subject to choose good objects, whether they appear sequentially or simultaneously.

**Fig. 4.**
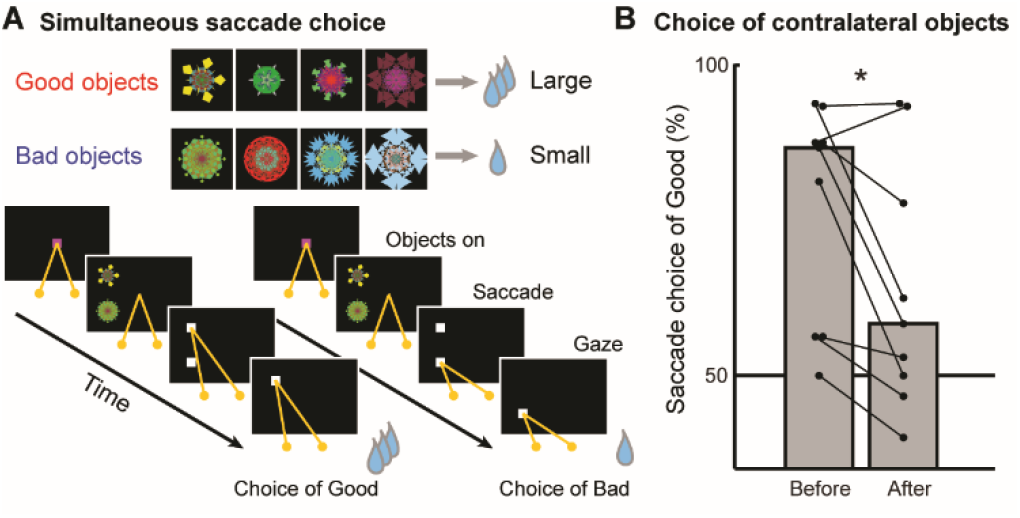
Blocking of the indirect pathway impairs choice of good objects. (A) Simultaneous saccade choice task. (B) Comparison of good object choice before vs. after the bicuculline injection in cvGPe (*n* = 9 sessions, \**P* = 0.027, Wilcoxon signed-rank test). Each bar indicates median of choice rate in contralateral choice. Each pair of connected dots depicts a single injection session.

## Discussion

Current data are consistent with that the direct pathway facilitates action (Go) and the indirect pathway suppresses action (No-go)(*18, 19*), while also consistent with that the direct and indirect pathways are active in actions(*20–22*) for the following reason. Because, when one action needs to be chosen (by the direct pathway), other actions must be suppressed (by the indirect pathway) at the same time. Overall, our data can evaluate these two contradicting theories by introducing the perspective of value-based action choice, and support the basal ganglia model that the indirect pathway suppresses unnecessary competing actions(*12, 14, 16*). This ‘rejection’ mechanism is important for visual search skill in humans and other animals. A recent study demonstrated that humans could quickly find a target object in visual search task, when they were instructed to ignore the specific feature of a distractor(*28*). The current study sheds light on the crucial role of the indirect pathway for rapidly finding a good object by ignoring bad objects (distractors) in periphery(*29*).

## Materials and Methods

### Subjects

Two adult male rhesus monkeys (*Macaca mulatta*, monkeys ZB and SP) were used for behavioral experiments, neuronal recordings and neuropharmacological experiments. All animal care and experimental procedures were approved by the National Eye Institute Animal Care and Use Committee and complied with the Public Health Service Policy on the Humane Care and Use of Laboratory Animals.

### Surgeries

A plastic head holder, plastic recording chambers, and scleral search coils were implanted under anesthesia with ketamine, diazepam, and isoflurane gas throughout these surgeries. After 6 weeks of recovery period, we started training and recording. We routinely cleaned the implants by flushing with hydrogen peroxide or a mixture of betadine and saline solution at least three times a week for the health and well-being of subjects.

### Behavioral procedures

Object-reward association learning task (Fig. 1A). This task was to let the subject learn the association of object with reward. This task procedure followed previous studies from the lab (*23, 24*). In each trial, one object (among 8 objects) was presented, to which the subject made a saccade. This was followed by a large or small reward, depending on the object (good or bad). The subject then learned the values of many objects (total 80 in ZB, total 48 in SP).

Passive viewing task (Fig. 1B). This task was used to investigate whether neurons encoded the values of individual objects more than one day after the object-reward association learning task. This task procedure followed the previous studies as well (*23, 24*). In each trial, several learned objects (1 – 6 objects) were chosen pseudo-randomly and presented sequentially at a peripheral position (i.e., neuron’s receptive field) while the subject was fixating at center. The trial ended with a reward, which however was not congruent with the presented objects. This task focused on the effect of object-value learning on neuronal responses before any behavioral response occurred.

Sequential saccade choice task (Fig. 2A, Movie S1 and S2). This task was used to investigate both neuronal and behavioral responses during and after object-value learning. A set of 8 objects were used in one block of 128 trials: 4 objects were associated with a large reward (good objects), while the other 4 objects were associated with no reward (bad objects) (Fig. 2A). In each trial, these objects were presented in a random sequence until the subject chose one (see below in detail). Each object appeared at one of four positions (up-right, down-right, up-left and down-left) pseudo-randomly (15° from center). After the subject fixated at a center green square for 700 ms, the fixation cue disappeared, and one object appeared. If the subject did not make a saccade to the object within 400 ms (i.e., no choice), another object appeared after the same fixating period (see 1st object in Fig. 2A). No choice occurred in another way: after making a saccade to the presented object within 400 ms, leave it within 400 ms, and return to the center (see 2nd object in Fig. 2A). In contrast, if the subject made a saccade to the object within 400 ms and then gazed at it for 400 ms (i.e., choice), the subjects received the outcome (reward or no reward) depending on the chosen object (good or bad). The next trials started after an inter-trial interval (1000 – 2000 ms). Overall, the subject learned to reject bad objects before choosing a good object to get the reward.

Simultaneous saccade choice task (Fig. 4A). This task was used to investigate whether the pharmacological manipulation modulated the learned choices between good and bad objects that appeared simultaneously at different positions. Before this task, the subject learned several sets (ZB: 3 sets and SP: 2 sets) of objects using the object-reward association learning task (Fig. 1A). One of them was used in one block of experiment including 64 choice trials. Each trial started with gaze fixation (700 ms) at a center magenta square. Then, the fixation cue disappeared and two randomly chosen objects (good and bad objects) appeared simultaneously, both in one hemifield (left or right) in the 50% of all trials, and in opposite hemifields (left and right) in the other 50%. When a saccade occurred (i.e., gaze 7° away from the fixation cue), these objects turned to white squares. After gazing at either square for 600 ms, a large or small reward was delivered depending on the subject’s choice (good or bad object).

### Neuronal recordings

Neuronal activity was recorded from two subjects (ZB and SP) in the caudal-ventral globus pallidus pars externus (cvGPe) and the caudal-dorsal-lateral substantia nigra pars reticulata (cdlSNr): 55 cvGPe neurons (ZB: 29 and SP: 26) and 52 cdlSNr neurons (ZB: 34 and SP: 18). The recording sites were determined with a 1mm spacing grid system, and MR images (4.7 T, Bruker) obtained along the direction of the chamber. Single-unit recording was performed using epoxy-coated tungsten microelectrodes (FHC). The electrodes were inserted into the brain through a stainless-steel guide tube and advanced by an oil-driven micromanipulator (MO-97A, Narishige). The electric signals from the electrode were amplified and band-bass filtered (0.2 – 10 kHz; BAK). All behavioral tasks and recordings were controlled by custom written visual C++ based software (Blip; http://www.robilis.com/blip/).

### Neuropharmacological manipulations

To temporarily block GABAergic inputs to cvGPe neurons, bicuculline (GABAA antagonist) was locally injected in cvGPe. The injection was done in the right cvGPe of each subject. We first recorded the neuronal activity encoding stable object values that are localized within cvGPe14, and then injected bicuculline. For this purpose, we used a custom-made injectrode consisting of a microelectrode (FHC) and a silica tube (Polymicro Technologies). Bicuculline methiodide (14343; Sigma-Aldrich) was dissolved in physiological saline. We injected 1 μl of 1 μg/μl bicuculline in each site at the speed of 0.2 μl/min using a manual pump. The subjects performed behavioral tasks starting 5 min after the injection. We studied the behavioral effect of bicuculline using the sequential saccade choice task (Fig. 2A) and the simultaneous saccade choice task (Fig. 4A). Main data were based on the comparison of saccades to (and from) the presented object between the pre-injection sessions and the post-injection sessions (within 1 hr after the injection). We sometimes studied the neuronal effect of bicuculline in cdlSNr that is the target of cvGPe (Fig. 3A and D). For control sessions, we injected 1 μl of physiological saline in each site in cvGPe at the speed of 0.2 μl/min. In total 9 sessions for bicuculline injections (ZB: 5 and SP: 4) and 8 sessions for saline injections (ZB: 5 and SP: 3) were conducted.

### Behavioral analysis

Saccade latency. Eye position was sampled at 1 kHz using a scleral search coil. Saccade latency was determined using software custom written with MATLAB (MathWorks). The saccade latency was measured from the object onset to the saccade onset.

Sequential saccade choice task. The main question of this task was whether a saccade followed by gaze occurred in response to the onset of an object. Such a saccade was detected when the eye position was out of the center window (10-degree square) within 400 ms from the object onset, while it was detected as no-saccade when the eye position stayed within the center window. After the saccade, it was judged as the acceptance of the object if the eye position stayed within the peripheral window (13-degree square around the object) for 400 ms, which was followed by an outcome (reward or no reward). Otherwise, the behavior was judged as the rejection of the object, as explained above (Behavioural procedures). The saccade rate (%) was calculated by (the number of trials in saccade to object) / (the total number of trials) × 100 (Fig. 2B, 3C and Fig. S3).

Simultaneous saccade choice task. The main question of this task was which object was chosen by the first saccade after two objects (good or bad) appeared simultaneously at different positions. The choice rate of good object (%) was calculated by (the number of trials in first saccade to good object) / (the total number of trials) × 100 (Fig. 4B and Fig. S4).

### Statistical analysis

Passive viewing task. To investigate whether cvGPe and cdlSNr neurons responded to bad objects, a paired t-test (two-tailed) was applied to neuronal spike counts. These responses to bad objects (from 100 ms to 300 ms after bad object onset) were compared with baseline activities (from 200 ms to 0 ms before bad object onset) in the same trials (Fig. 1C and D).

Sequential saccade choice task. To investigate the difference in behavioral (saccade) response to good and bad objects, we used a Mann-Whitney *U* test (two-tailed) for saccade latency (Fig. S1) and for saccade choice more than month after the last session of the task (Fig. 2B). To investigate the neuronal data, we used a paired t-test (two-tailed) (Fig. 2C and D). Total 55 cvGPe neurons and 52 cdlSNr neurons were used for this analysis, respectively. To investigate whether these cvGPe and cdlSNr neurons responded to bad objects, their responses to bad objects (from 100 ms to 300 ms after bad object onset) were compared with baseline activities (from 200 ms to 0 ms before bad object onset) in the same trials. To investigate whether these cvGPe and cdlSNr neurons showed different response depending on action choice to bad objects (saccade or not), their activity was compared between saccade (+) trials and saccade (-) trials in the same spike count window (from 100 ms to 200 ms after bad object onset).

Neuropharmacological experiments. To investigate whether the bicuculline injection affected cdlSNr activity in the sequential saccade choice task, a Mann-Whitney *U* test (two-tailed) was applied for comparing the activity of the same cdlSNr neuron between before and after the injection, separately for good or bad objects (Fig. 3D). To investigate whether the bicuculline injection modulated saccade rate to bad objects in the sequential saccade choice task, a Wilcoxon signed-rank test (two-tailed) was applied for comparing the saccade rates between before and after bicuculline injection (Fig. 3C and Fig. S3A) and saline injection (Fig. S3B). To investigate whether the injection modulated the saccade choice rate to good objects in the simultaneous saccade choice task, a Wilcoxon signed-rank test (two-tailed) was applied for comparing the saccade choice rates of good objects between before and after bicuculline injection (Fig. 4B and Fig. S4A) and saline injection (Fig. S4B).

## Acknowledgments

We thank M. Smith for creating injectrodes, A.M. Nichols for creating chambers, D. Parker for help with animal care and surgery, and A. Ghazizadeh, H. F. Kim, M. Isoda, and Y. Tachibana for technical advice, and J. Kunimatsu, K. Maeda, D. McMahon, G. Atul, G. Costello, H. Lee, A. Yoshida, J.P. Herman and B. Amarender for comments.

## Author contributions

H.A. and O.H. conceived and designed experiments. H.A. performed all experiments. H.A. and O.H. formed the conceptual model. H.A. analyzed the data and made all figures. H.A. and O.H. wrote the paper.

## Data and materials availability

The data analyzed, and computer code used for data analysis are available from the corresponding author on reasonable request.

