## Supplemental Figure S1 for "Indirect pathway of caudate tail for choosing good objects in periphery"

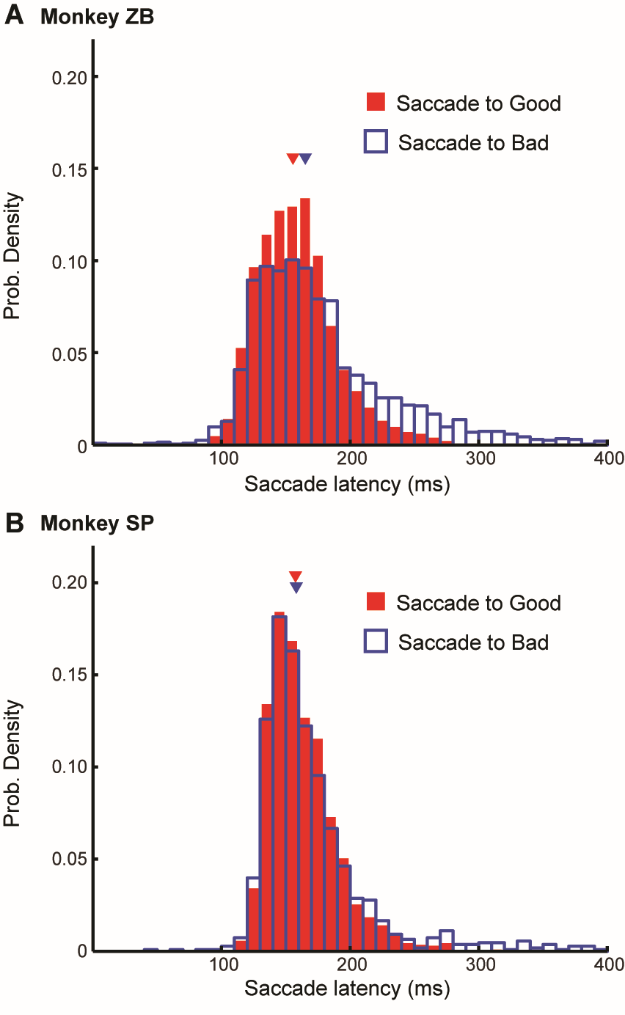
**Fig. S1.**

**Latency of saccade to good and bad objects in sequential saccade choice task.** (A) Probability density functions of saccade latencies in Monkey ZB to good objects (red) and bad objects (blue) (Good; *n* = 3934 trials, Bad; *n* = 2033 trials, ****P* < 0.001, Mann-Whitney *U* test). (B) Same data in Monkey SP (Good; *n* = 2281 trials, Bad; *n* = 1080 trials, *P* = 0.12, Mann-Whitney *U* test).
