## Supplemental Figure S2 for "Indirect pathway of caudate tail for choosing good objects in periphery"

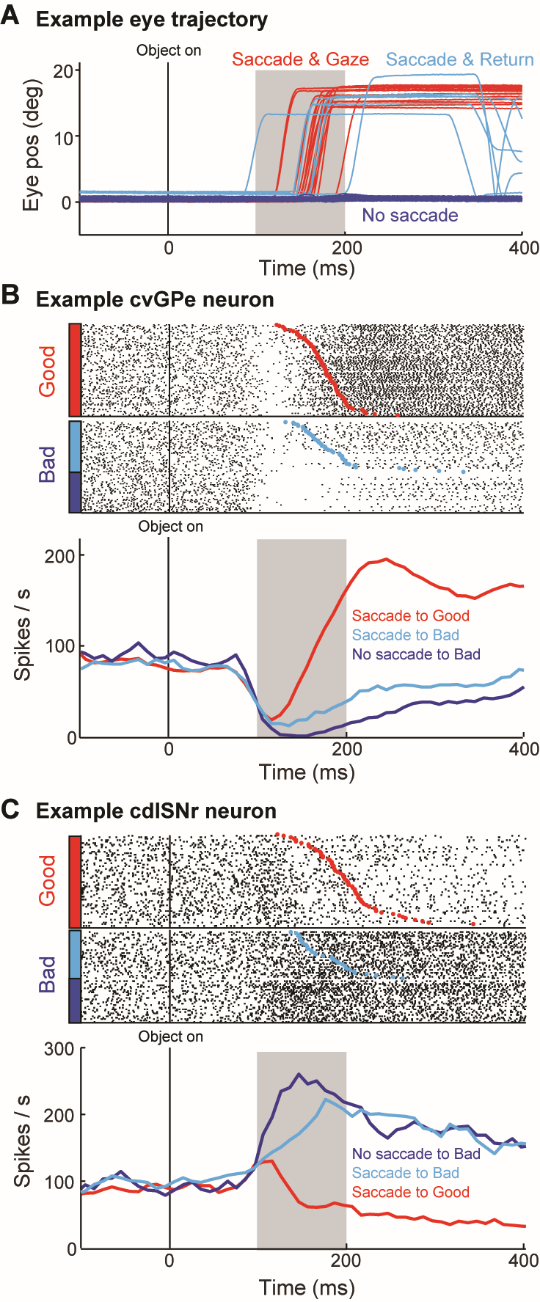
**Fig. S2.**

**Saccades and neuronal activities during sequential saccade choice task.** (A) Eye positions (eccentricity from center) aligned by object onset, showing saccades to good objects followed by sustained gaze (red), saccades to bad objects followed by returning saccades (cyan), and no saccade to bad objects (blue). (B) Activity of a cvGPe neuron. Top: Raster plots (same format as in Fig. 1C). Red and cyan dots indicate the onsets of saccades to good and bad objects, respectively. Bottom: Spike density functions (same format as in Fig. 2C). (C) Activity of a cdlSNr neuron.
