## Supplemental Figure S3 for "Indirect pathway of caudate tail for choosing good objects in periphery"

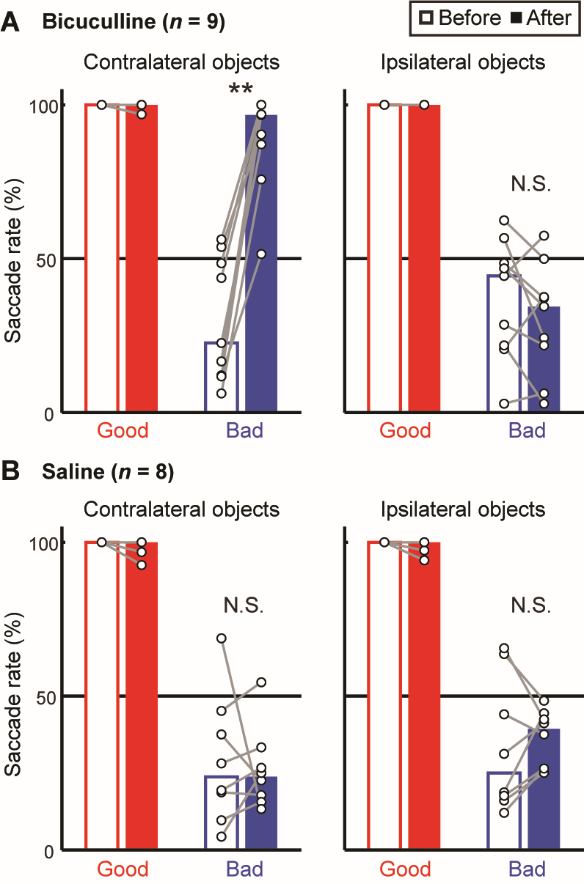
**Fig. S3.**

**Selective change in saccade rate by cvGPe manipulation during sequential saccade choice task.** (A) Effects of bicuculline injection in cvGPe (same format as in Fig. 3C). (B) Effects of saline injection in cvGPe (same format as in Fig. 3C). The only significant change by injection occurred in saccades to contralateral bad objects after the bicuculline injections (A, left; same data as Fig. 3C); saccades to ipsilateral bad objects were unchanged (A, right; *n* = 9 sessions, *P* = 0.34, Wilcoxon singed-rank test). The saline injection did not affect saccade rate to either contralateral bad objects (B, left; *n* = 8 sessions, *P* = 0.74, Wilcoxon singed-rank test) or ipsilateral bad objects (B, right; *n* = 8 sessions, *P* = 0.64, Wilcoxon singed-rank test).
