## Supplemental Figure S4 for "Indirect pathway of caudate tail for choosing good objects in periphery"

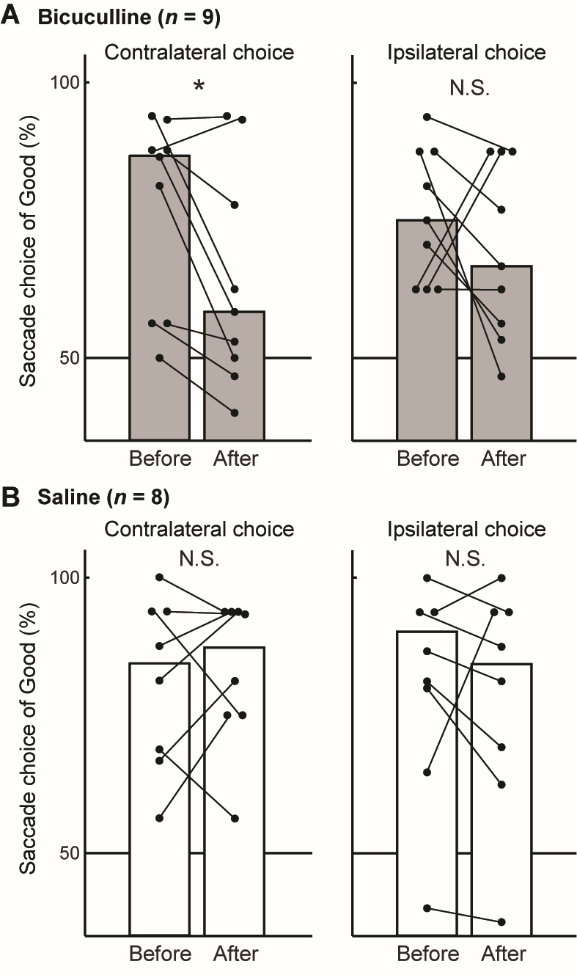
**Fig. S4.**

**Selective change in choice rate by cvGPe manipulation during simultaneous saccade choice task.** (A) Effects of bicuculline injection in cvGPe (same format as in Fig. 4B). (B) Effects of saline injection in cvGPe (same format as in Fig. 4B). The only significant change by injection occurred in choice of contralateral good objects after bicuculline injection (A, left; same data as Fig. 4B); choice of ipsilateral good objects was not significantly changed (A, right; *n* = 9 sessions, *P* = 0.52, Wilcoxon signed-rank test). Saline injection did not significantly affect choice of good objects in either contralateral choice (B, left; *n* = 8 sessions, *P* = 0.92, Wilcoxon signed-rank test) or ipsilateral choice (B, right; *n* = 8 sessions, *P* = 0.63, Wilcoxon signed-rank test).
