## Supplemental Movie Legends for "Indirect pathway of caudate tail for choosing good objects in periphery"

Movie S1.

**An example trial of sequential saccade choice task with eye movements.** See Fig. 2A. Yellow dot, subject’s eye position. Center green square, fixation cue. Each fractal object sequentially appears in periphery. The text “Bad” or “Good” appears in the upper right corner, when the bad or good object is presented (This text was not shown in actual experiments). In this trial, the subject rejected all bad objects, then accepted the final good object followed by reward delivery. The video is slowed down 3 times.

Movie S2.

**Example trials of sequential saccade choice task.** Same convention as for Movie S1. The subject rejected all bad objects, then chose good objects by gaze. The movie plays at real-time speed.

Movie S3.

**Example trials of sequential saccade choice task after bicuculline injection.** Same convention as for Movie S1. Bicuculline injection site was right cvGPe. The subject frequently made saccade to bad objects at left hemifield (contralateral side), but not right hemifield (ipsilateral side). The movie plays at real-time speed.

Movie S4.

**Example trials of sequential saccade choice task after saline injection.** Same convention as for Movie S1. Saline injection site was right cvGPe. The subject often ignored bad objects at both hemifields, then chose good objects followed by reward delivery. The movie plays at real-time speed.
